# A MinD-like ATPase couples flagellation and cell division in spirochetes

**DOI:** 10.64898/2026.04.08.717139

**Authors:** Kai Zhang, Wangbiao Guo, Michael J. Lynch, Brian R. Crane, Jun Liu, Chunhao Li

## Abstract

Spirochetes are evolutionarily distinct bacteria defined by their spiral morphology, unique means of motility, and periplasmic flagella (PFs). Because these filaments reside within the periplasm and are mechanically integrated with the cell body, their assembly must be precisely coordinated with cell growth and cytokinesis. However, the mechanism that couples flagellar biogenesis to cell division in spirochetes remains unclear. Using the Lyme disease spirochete *Borrelia burgdorferi* as a model, we identify FlhG (BB0269), a MinD-like ATPase, as a spatial regulator that links cell division to flagellar patterning. In wild-type cells, 7-11 long helical PFs originate from cell poles and assemble into ribbon-like bundles that wrap around the cell cylinder to drive motility. Deletion of *flhG* disrupts this ordered architecture, causing marked heterogeneity in flagellar number, defective ribbon assembly, aberrant septation, and severe motility impairment. Mechanistically, FlhG dynamically localizes to the poles and midcell during division, where it directs the positioning of FlhF, a signal recognition particle (SRP) -type GTPase controlling flagellar number and placement, and FliF, the MS-ring protein that nucleates flagellar assembly. Through this spatial regulation, FlhG coordinates flagellar assembly with cytokinetic progression. Together, these findings reveal a spatial regulatory mechanism coupling cell division to flagellation, providing insight into understanding how spirochetes coordinate their distinctive morphogenesis, flagellation and motility.

**Significance:** Spirochetes such as *Borrelia burgdorferi*, the causative agent of Lyme disease, rely on periplasmic flagella for motility and cell shape, yet how these structures are coordinated with cell division has remained unclear. We identify a MinD-like ATPase, FlhG, as a spatial regulator that couples flagellar assembly to cytokinesis. In contrast to its homologs in other bacteria, FlhG does not regulate flagellar protein levels but instead directs subcellular positioning of key assembly factors. By dynamically redistributing between the cell poles and division site, FlhG synchronizes flagellar patterning with septum formation. These findings uncover a previously unrecognized mechanism linking cell morphogenesis to the cell cycle and reveal how conserved ATPases can be repurposed to organize complex bacterial architectures.

## INTRODUCTION

Spirochetes represent a deeply branching lineage of bacteria distinguished by their spiral morphology and periplasmic flagella (PFs) (1–3). This phylum includes major human pathogens such as *Borrelia burgdorferi* (Lyme disease) (4), *Treponema pallidum* (syphilis) (5), and *Leptospira interrogans* (leptospirosis) (6). Unlike the external flagellate model organisms such as *Escherichia coli* and *Bacillus subtilis*, spirochetal PFs are confined within the periplasmic space, where they originate at both poles and extend inward along the cell cylinder (1). In most spirochetal species, these filaments assemble into ribbon-like bundles that wrap around the cell body, generating both the characteristic helical shape and the undulatory or corkscrew-like motility essential for environmental adaptation and host colonization (1, 2, 7-11). Thus, in spirochetes, flagella are structural determinants of cell morphology as well as engines of motility (1, 2, 12).

The integration of PFs within the periplasm imposes unique spatial constraints on their assembly (1, 2). During cell elongation and division, new poles are generated and daughter cells must establish the correct number, placement, and organization of PFs to preserve morphology and motility. Failure to properly pattern these filaments disrupts ribbon architecture, alters cell shape, and compromises locomotion (1, 13–15). In bacteria with polar flagella, the number and positioning of flagella are controlled by conserved regulators such as FlhF and FlhG (16–18); whether and how these systems are integrated with the division machinery in spirochetes remains unknown.

More broadly, this gap reflects a fundamental question in bacterial cell biology: how are large macromolecular machines spatially coordinated with the cell cycle to maintain cellular architecture? The structural role of PFs suggests that spirochetes require mechanisms that directly couple cytokinesis to flagellar biogenesis, yet such mechanisms have not been defined. Here, using *B. burgdorferi* as a model system, we uncover a spatial regulatory mechanism that links cytokinesis to flagellar patterning, revealing how spirochetes coordinate morphogenesis and flagellation to maintain their distinctive cellular organization and unusual means of motility.

## RESULTS

### BB0269 is a MinD-like ATPase

BB0269 (295 amino acids; predicted molecular mass 32.8 kDa) is located within the *flgB* motility gene operon and is annotated as a FlhG-like ATP-binding protein (19, 20) (hereafter referred to as FlhG_Bb_). Sequence analysis revealed that FlhG_Bb_ belongs to the MinD/ ParA family of P-loop NTPases (16, 21), sharing 25.73% and 32.58% sequence identity with *E. coli* MinD (22) and *Pseudomonas aeruginosa* FleN (a FlhG homolog) (23), respectively. Similar to other members of this family (16, 24), FlhG_Bb_ contains three conserved ATPase motifs: a deviant Walker A motif or P-loop (³⁸GKGGVGKS⁴⁵), Switch I (⁶⁷DADIGMAN⁷⁴), and Switch II (¹⁴⁷DTSAGIS¹⁵³) (**Fig. 1A** and **Fig. S1 and 2**).

**Figure 1.**
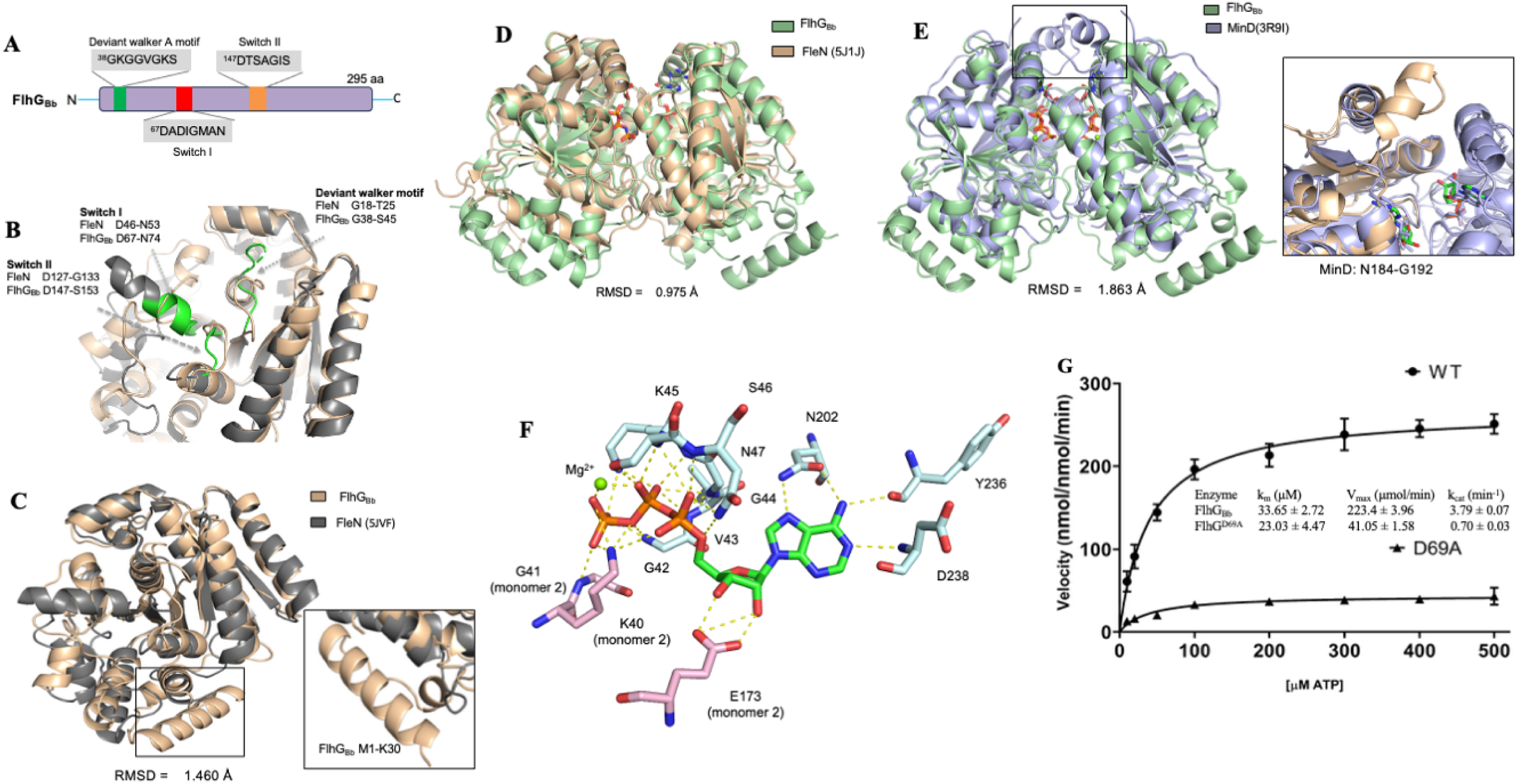
Structural and biochemical analyses of FlhG (FlhG_Bb_) from *B. burgdorferi*. **(A)** Schematic illustration of FlhG_Bb_ domain organization. The positions and amino acid sequence of the deviant Walker A, switch I, and switch II motifs are indicated. **(B)** Structural comparison of FlhG_Bb_ and *P. aeruginosa* FleN (PDB: 5JVF) subunits shows the deviant Walker A, switch I, and switch II motif labeled in green. **(C)** Structural superimposition of FlhG_Bb_ (tan) with FleN (PDB: 5JVF; dark gray) identifies a unique N-terminal α-helical extension (inset: M1-K30). **(D)** Structural superimposition of FlhG_Bb_ (green) with FleN (PDB: 5J1J; tan) dimers. RMSD = 0.975 Å. **(E)** Structural superimposition of the FlhG_Bb_ dimer (green) with the MinD dimer (PDB: 3R9I; blue). Inset shows a difference in the position of an interfacial helix comprising residues N184-G192. **(F)** Close-up view of the ATP-binding active site of FlhG_Bb_ showing key residues from both monomers involved in nucleotide coordination. **(G)** Michaelis-Menten kinetics of ATPase activity for wild-type FlhG_Bb_ (WT) and D69 point mutant (D69A). Kinetic parameters (K_m_, V_max_, and k_cat_) are shown in the inset table. Data represent mean ± SD.

The AlphaFold structural model of FlhG_Bb_ closely resembles both MinD (RMSD = 1.863 Å; PDB: 3R9I (25)) and FleN (RMSD = 1.460 Å; PDB: 5JVF (26)) (**Fig. 1B**). FlhG_Bb_ displays greater structural similarity to FleN than to MinD. However, structural modeling suggests that, unlike FleN, FlhG_Bb_ does not undergo a major conformational change upon ATP binding, either in the monomeric (**Fig. S3**) or dimeric states (**Fig. S4**). The other notable difference is that FlhG_Bb_ contains a unique ∼30-residue N-terminal α-helical extension (residues M1-S30) that is absent from both MinD and FleN (**Fig. 1C**). Structural superimposition further predicts that FlhG_Bb_ adopts a dimeric structure similar to that of *E. coli* MinD. However, the model reveals a distinct repositioning of an interfacial helix (residues N184-G192) at the predicted dimer interface (**Fig. 1D**), a configuration that more closely resembles that observed in FleN (**Fig. 1E**). Despite these structural variations, FlhG_Bb_ retains a conserved catalytic pocket predicted to coordinate ATP-Mg²⁺ binding (**Fig. 1F**).

To determine whether FlhG_Bb_ is enzymatically active, we performed in vitro ATPase assays using purified recombinant proteins. FlhG_Bb_ exhibited robust ATP hydrolysis activity following single-substrate steady-state kinetics (**Fig. 1G**). Substitution of Asp69 with alanine (D69A), a conserved residue required for ATP hydrolysis (26, 27), nearly abolished ATPase activity. Collectively, these findings demonstrate that FlhG_Bb_ functions as a bona fide ATPase.

### Deletion of *flhG_Bb_* impairs *B. burgdorferi* growth and alters cell morphology

The *flhG_Bb_* gene is located between *flhF* (*BB0270*)(13) and *flgV* (*BB0268*) (28, 29) in the *flgB* motility gene operon (20). To minimize potential polar effects on the expression of adjacent genes, *flhG_Bb_* (*BB0269*) was deleted and replaced in-frame with a streptomycin-resistance cassette (*aadA1*) (**Fig. 2A**). The resulting mutant (Δ*flhG*) was complemented by inserting the *flgB-BB0269* fragment into the intergenic region between *BB0445* and *BB0446*, as previously described (30) (**Fig. 2B**). Immunoblot analysis confirmed that FlhG_Bb_ was absent in the Δ*flhG* mutant and restored in the complemented strain (Δ*flhG*^com^) (**Fig. 2C**). The Δ*flhG* mutant exhibited a pronounced growth defect (**Fig. 2D**). At stationary phase, the mutant reached a cell density of 2.34 × 10⁷ cells ml⁻¹, representing an approximately 7-fold reduction compared with the wild type (1.59 × 10⁸ cells ml⁻¹). This growth defect was fully rescued by complementation.

**Figure 2.**
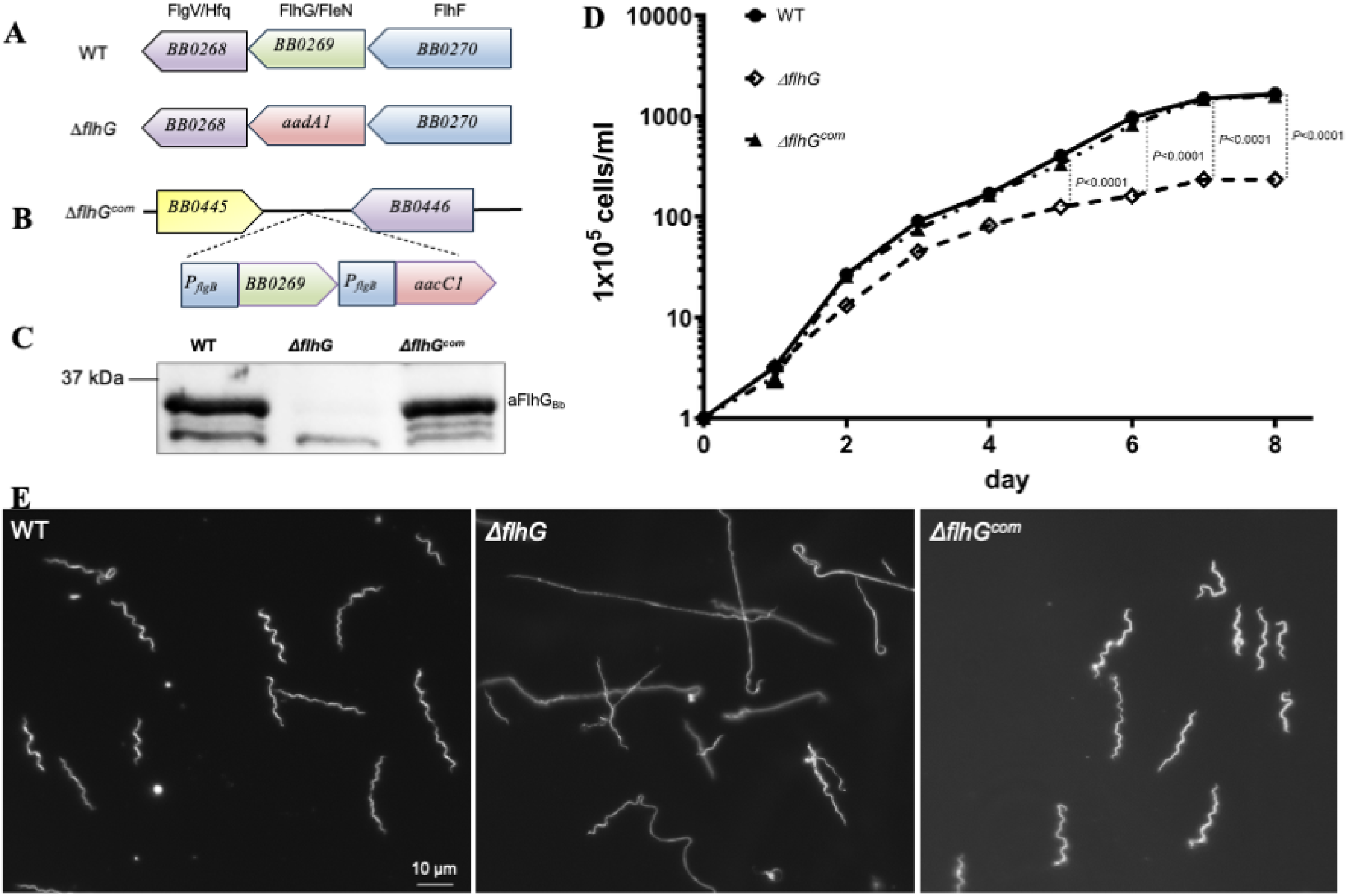
FlhG_Bb_ is required *for B. burgdorferi* normal growth and morphology. **(A)** Schematic of the chromosomal gene locus surrounding *flhG* (*BB0269*) in wild-type (WT) and the *ΔflhG* deletion mutant, in which *flhG* is in-frame replaced by the *aadA1* spectinomycin resistance cassette. **(B)** Schematic of cis-complementing the *ΔflhG* mutant by inserting *flgB-BB0269* into the intergenic region between BB0445 and BB0446, alongside a gentamicin resistance cassette (*aacC1*). **(C)** Immunoblot confirming FlhG_Bb_ expression in WT, *ΔflhG*, and *ΔflhG*^com^ strains probed with anti-FlhG_Bb_ antibody (αFlhG_Bb_). **(D)** Growth curves of WT, *ΔflhG*, and *ΔflhG*^com^ strains in BSK-II medium. Cell density was measured for up to 8 days using a Petroff–Hausser counting chamber. Statistical comparisons were performed at days 6, 7, and 8 (*P < 0.0001*). **(E)** Dark-field microscopy images of WT, *ΔflhG*, and *ΔflhG*^com^ strains. Images were taken at later-log growth phase. Scale bar = 10 μm.

Dark-field microscopy further revealed striking morphological abnormalities in mutant cells. Wild-type cells displayed the characteristic wave-like morphology and were relatively uniform in length. In contrast, most Δ*flhG* cells adopted a rod-like morphology and frequently formed chains (**Fig. 2E**). Detailed measurements revealed two predominant mutant populations: elongated rod-shaped cells (>70% of cells; average length 39.41 ± 13.33 μm, *n* = 22) and short spiral cells (<30%; average length 11.53 ± 3.48 μm, *n* = 16). The elongated rods were approximately twice the length of wild-type cells (17.27 ± 3.39 μm, *n* = 37), whereas the short cells were roughly half the wild-type length. These results indicate that loss of FlhG_Bb_ disrupts normal cell division and morphogenesis in *B. burgdorferi*.

### FlhG_Bb_ is essential for the motility of *B. burgdorferi*

To assess the role of FlhG_Bb_ in motility, we performed swimming plate assays in 0.35% semisolid agar. The Δ*flhG* mutant produced significantly smaller migration rings than the wild type (WT: ∼15.2 mm vs. Δ*flhG*: ∼7.9 mm; *P* < 0.0001) (**Fig. 3A, B**). Complementation with wild-type *flhG_Bb_* fully restored the migration ring diameter (∼15.2 mm). In contrast, complementation with an ATPase-deficient point mutant (Δ*flhG^D69A^*) failed to rescue motility (∼7.6 mm; *P* < 0.0001 vs. WT) (**Fig. 3B**). These results indicate that the ATPase activity of FlhG_Bb_ is required for normal motility. Consistent with the swimming plate assays, quantitative tracking analysis revealed that wild-type and complemented (Δ*flhG*^com^) cells displayed robust translational movement (**Videos 1 and 3**) and their averaged velocity in 1% methylcellulose reached ∼ 11 μm/s (WT: 75 cells; Δ*flhG*^com^: 96 cells), whereas Δ*flhG* and Δ*flhG^D69A^*cells exhibited wave-like body undulation (**Videos 2 and 4**) but failed to move (∼ 0 μm/s; Δ*flhG:* 47 cells; Δ*flhG^D69A^*: 71 cells) (**Fig. 3C**). Together, these findings demonstrate that FlhG_Bb_ and its ATPase activity are essential for *B. burgdorferi* motility.

**Figure 3.**
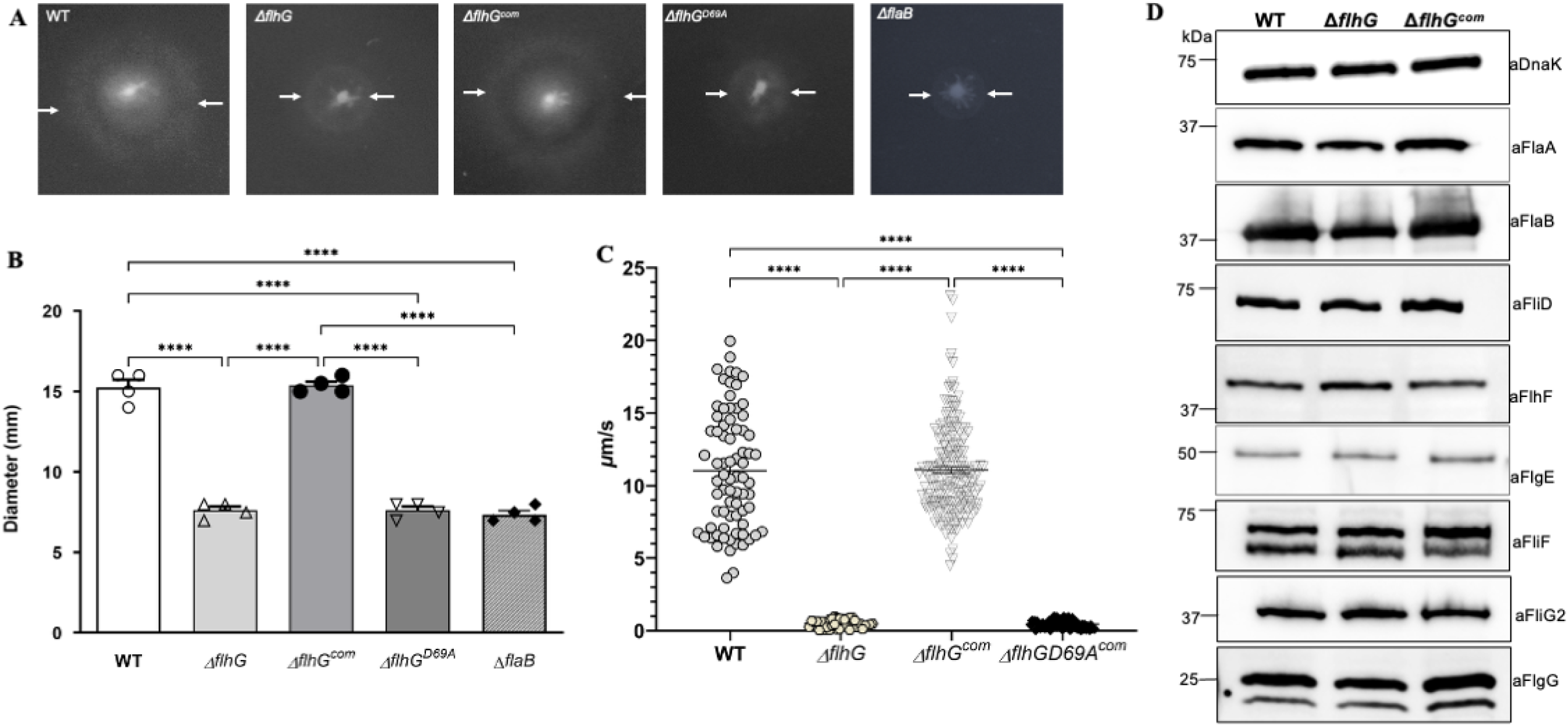
FlhG ATPase activity is required for motility but not flagellar protein synthesis. **(A)** Simming plate assays of WT, *ΔflhG*, *ΔflhG*^com^, and *ΔflhG*^D69A^ strains. *ΔflaB*, a non-motile mutant(12), was used as a control to measure the inoculum size. Arrow pairs indicate the outer border of swimming rings. **(B)** Quantification of swimming ring diameters (mm) for the indicated strains. Data shown as mean ± SD with individual data points. **** *P < 0.0001*. **(C)** Swimming speeds (μm/s) of individual cells for WT, *ΔflhG*, *ΔflhG*^com^, and *ΔflhG^D69A^*strains measured by bacterial motion tracking analysis. **** *P < 0.0001*. **(D)** Immunoblot analysis of flagellar and motor structural proteins in WT, *ΔflhG*, and *ΔflhG*^com^ strains. Whole-cell lysates were probed with antibodies against DnaK (loading control), FlaA, FlaB, FliD, FlhF, FlgE, FliF, FliG2, and FlgG as indicated. Molecular weight markers (kDa) are shown.

### FlhG_Bb_ does not regulate flagellar protein synthesis

FlhG is generally considered as a negative regulator of flagellar number; accordingly, deletion of *flhG* in many bacteria leads to elevated levels of flagellar proteins and aberrant flagellation (16, 31–34). To determine whether this regulatory effect occurs in *B. burgdorferi*, we compared the abundance of several representative flagellar proteins in wild-type and Δ*flhG* strains by immunoblotting. The proteins examined included the MS-ring protein FliF(35), the motor switch protein FliG2 (8), the major flagellin FlaB (12), the flagellar sheath protein FlaA (36), the filament cap protein FliD (37), and the SRP-like GTPase FlhF (13). Unexpectedly, all tested proteins were present at comparable levels in both strains (**Fig. 3D**), indicating that the canonical role of FlhG in controlling flagellar protein abundance is not conserved in *B. burgdorferi*. Thus, unlike its homologs in other bacteria (32–34, 38), FlhG_Bb_ appears to govern motility independently of flagellar protein production, implicating alternative downstream mechanisms in its function.

### Loss of FlhG_Bb_ leads to aberrant flagellar patterning

FlhG_Bb_ is essential for *B. burgdorferi* motility; however, its deletion does not significantly affect flagellar protein synthesis, leaving the mechanistic basis of the motility defect unclear. To address this, we examined flagellar architecture in Δ*flhG* and complemented strains by cryo-electron tomography (cryo-ET). Previous studies have shown that *B. burgdorferi* cells typically have 7-11 PFs per pole, which extend inward and form two ribbon-like bundles that tightly wrap around the cell cylinder (7, 13, 39). In Δ*flhG* cells, however, this pattern was severely disrupted: PF numbers were highly variable, ranging from 2 to 19 per cell pole, and flagellar motors were distributed without discernible pattern, failing to coalesce into ribbon-like structures (**Fig. 4A, B, D**). A comparable disorganization was observed in the D69A point mutant (**Fig. 4E**), indicating that the regulatory function of FlhG_Bb_ on flagellar patterning depends on its ATPase activity. In contrast, complemented cells fully restored the wild-type flagellar pattern, with 7-11 PFs forming characteristic tight ribbon bundles that wrap around the cell body and extend toward the cell center (**Fig. 4C, F**). Collectively, these findings establish that FlhG_Bb_ governs both flagellar number and spatial organization, two properties that are essential for coordinated motility in *B. burgdorferi*.

**Figure 4.**
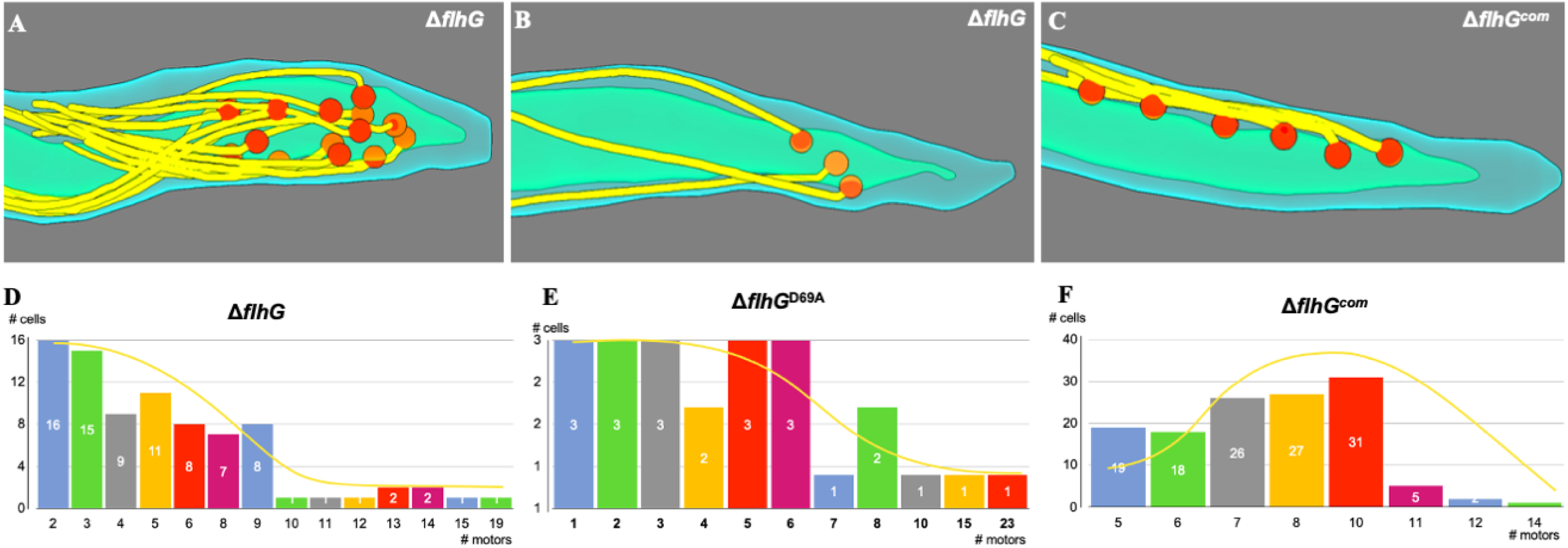
Loss of *flhG_Bb_* results in dysregulation of flagellar motor number. **(A-C)** Representative three-dimensional segmentation of tomograms illustrating flagellar motor distributions in *ΔflhG* cells with many motors (A), few motors (B), and a *ΔflhG*^com^ cell with restored motor number (C). **(D-F)** Histograms showing the distribution of flagellar motor numbers per cell for *ΔflhG* (D), *ΔflhG*^D69A^ (E), and *ΔflhG*^com^ (F). Numbers inside bars indicate cell counts. The yellow curves represent expected Poisson distributions.

### Loss of FlhG_Bb_ disrupts the flagellar motor collar structure

In addition to flagellar patterning, we also examined the potential impact of FlhG_Bb_ on the ultrastructure of flagellar motors *in situ* using cryo-ET and subtomogram averaging. Strikingly, subtomogram averaging revealed that a ring-like structure present in the wild-type flagellar motor (**Fig. 5A & D**) was absent in the *ΔflhG* mutant (**Fig. 5B & E**). Importantly, this structural defect was fully rescued in the complemented strain (**Fig. 5C & F**). This ring-like structure sits atop the flagellar collar, a hallmark feature unique to spirochetal flagellar motors (40–42), suggesting that FlhG_Bb_ plays a role in motor architecture. Collectively, these findings establish FlhG_Bb_ as a critical determinant of collar integrity, though the precise molecular mechanism by which it governs flagellar collar assembly remains to be determined.

**Figure 5.**
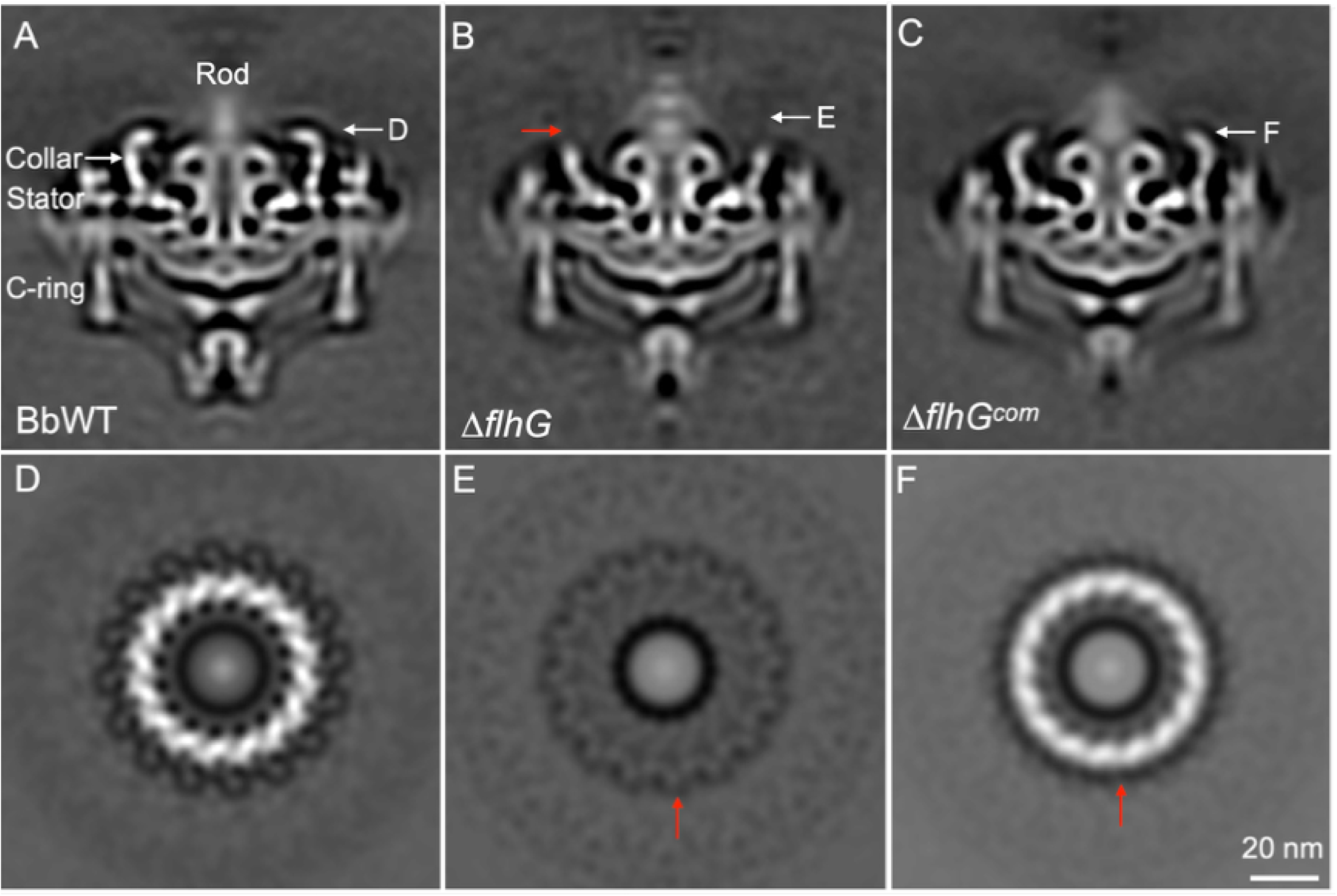
FlhG_Bb_ is required for proper flagellar collar assembly as revealed by cryo-ET. **(A-C)** Longitudinal cross-sectional views of subtomogram averages of the flagellar motor from WT (A), *ΔflhG* (B), and *ΔflhG*^com^ (C) strains. Arrows indicate the positions of the disk (D-F) structures referenced in panels D-FA-C. **(D-F)** Top-down cross-sectional views at the outer disk for WT (D), *ΔflhG* (E), and *ΔflhG*^com^ (F). Red arrows indicate the disk density. Deletion of *flhG_Bb_* results in loss of the characteristic disk density (E), which is restored upon complementation (F).

### Deletion of *flhG_Bb_* leads to aberrant cell division

Deletion of *flhG_Bb_* affects *B. burgdorferi* growth and cell morphology (**Fig. 2**), suggesting a potential role in cell division. To test this possibility, we employed hydroxycoumarin–D-alanine (HADA) labeling (43) to visualize newly synthesized peptidoglycan in live cells and monitor septum formation. Consistent with a previous report (44), wild-type cells exhibited discrete HADA signals at division sites that followed the characteristic 1/4, 1/2, and 3/4 pattern (**Fig. 6A**). Demographic analysis further showed that most wild-type cells displayed a single HADA band at midcell (**Fig. 6B,** n=936 cells). In contrast, this spatial pattern was markedly disrupted in Δ*flhG* cells (**Fig. 6C, D,** n=900 cells). Specifically, 66.7 ± 7.3% of mutant cells exhibited irregular high-intensity HADA signals that did not conform to the 1/4, 1/2, and 3/4 rule for the fraction of the cell outline being fluorescently marked at the early, mid and late stages of growth. In addition, 20.9 ± 6.5% of cells displayed strong signals at one cell end, 7.4 ± 3.8% at midcell, and 4.9 ± 3.8% at both cell ends. These aberrant labeling patterns indicate that septum positioning and peptidoglycan synthesis become spatially misregulated in the absence of FlhG. Together, these findings demonstrate that FlhG_Bb_ contributes to proper division-site placement in *B. burgdorferi*, supporting a role for this ATPase in coordinating cell division with flagellar organization.

**Figure 6.**
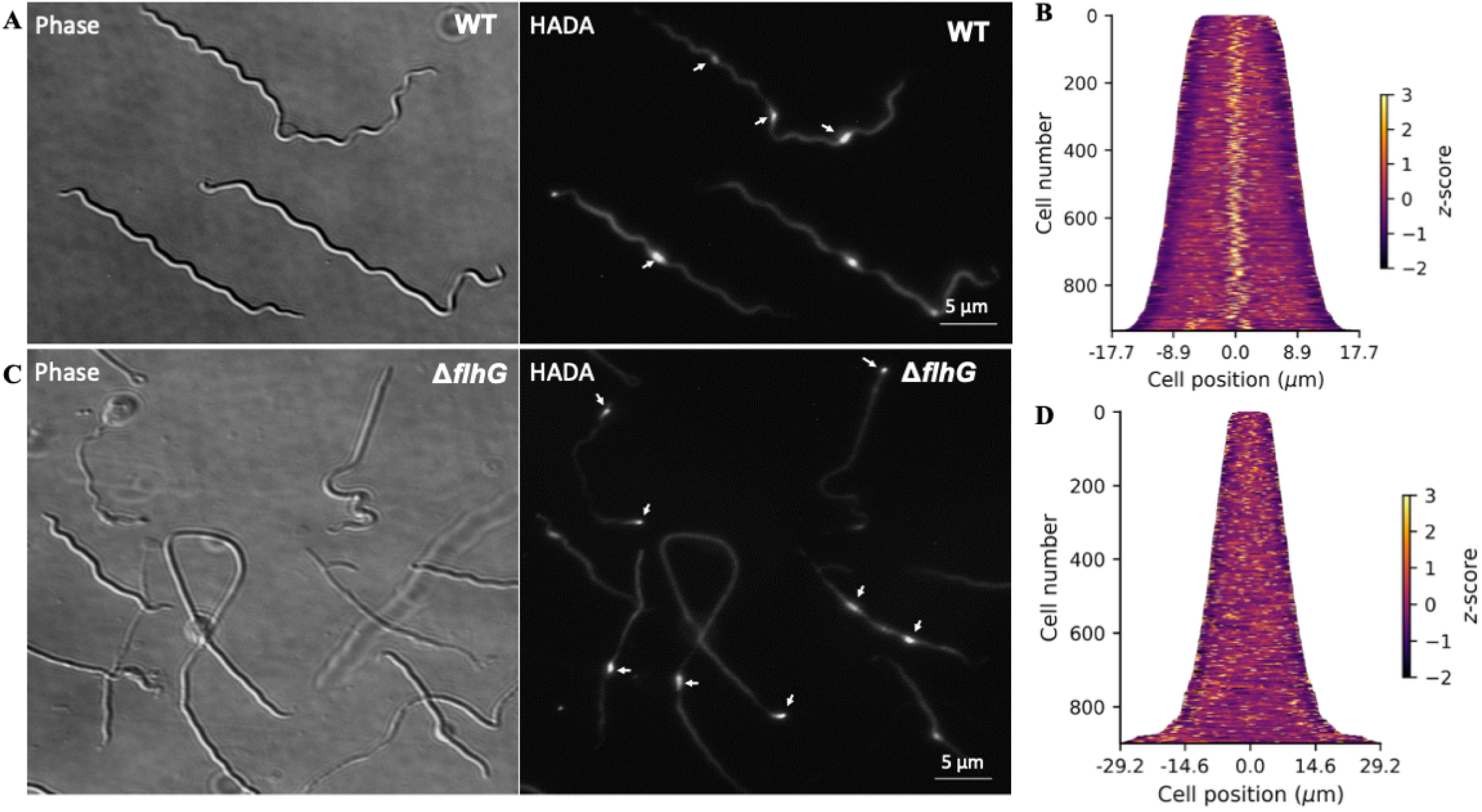
Loss of flhG_Bb_ leads to aberrant cell division in *B. burgdorferi*. **(A, C)** Phase contrast (left) and HADA fluorescence (right) images of WT (A) and *ΔflhG* (C) cells labeled with the fluorescent D-amino acid HADA to visualize sites of active peptidoglycan synthesis. White arrows indicate HADA incorporation foci. Scale bar = 5 μm. **(B, D)** Heat maps of normalized HADA fluorescence intensity as a function of cell body position (μm) for WT (B) and *ΔflhG* (D). Each row represents a single cell (n ≥ 900 cells per strain), with z-score-normalized fluorescence depicted on a color scale (purple to yellow). *ΔflhG* cells show a wider distribution of cell lengths compared to WT.

### FlhG_Bb_ localizes to the cell poles and midcell

To determine how FlhG_Bb_ regulates flagellar patterning and cell division, we examined its subcellular localization using a GFP fusion approach. A plasmid expressing FlhG_Bb_-GFP was constructed and introduced into the *flhG* mutant strain. Immunoblot analysis with antibodies against GFP and FlhG_Bb_ confirmed expression of the fusion protein (**Fig. S5A**). Fluorescence microscopy (**Fig. 7A**) combined with demographic analysis (**Fig. 7B**) revealed that GFP puncta were predominantly detected at the cell poles in short newborn cells, whereas in longer dividing cells fluorescence signals were observed at both poles and the midcell division site. A similar localization pattern was observed when the FlhG_Bb_-GFP expression vector was introduced into wild-type cells (**Fig. S5B**). Together, these results indicate that FlhG_Bb_ exhibits dynamic localization, residing primarily at the cell poles in nascent cells and redistributing to the midcell division zone during cell elongation and division. This dynamic, cell-cycle-dependent localization, reminiscent of MinD in rod-shaped bacteria (45), likely directs proper flagellar assembly and synchronizes it with cytokinesis in *B. burgdorferi*.

**Figure 7.**
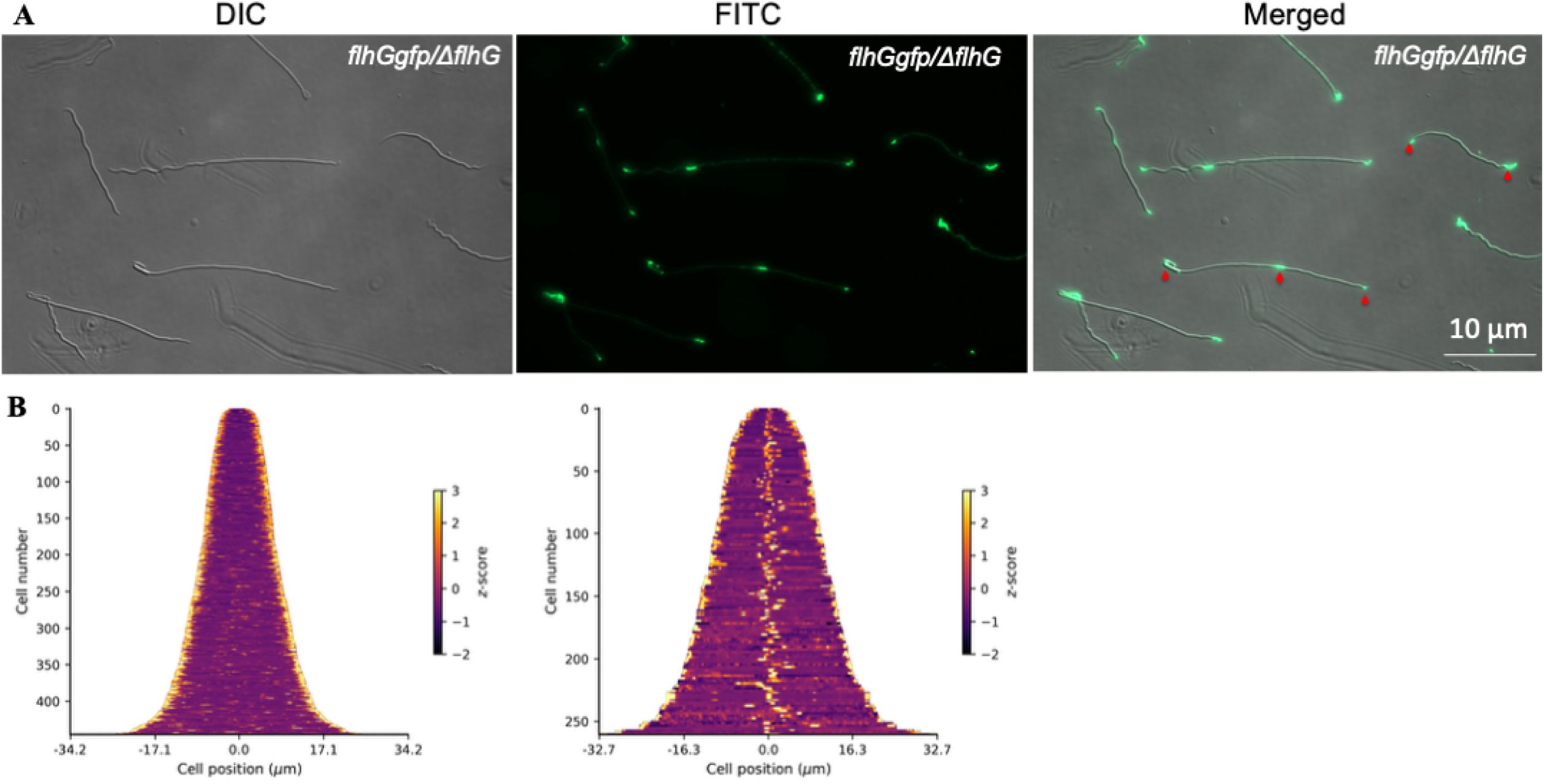
FlhG-GFP localizes to the flagellar insertion points in *B. burgdorferi*. (A) Fluorescence microscopy of *flhGgfp/ΔflhG* cells expressing FlhG-GFP as the sole copy of FlhG. DIC (left), FITC fluorescence (middle), and merged (right) channels are shown. FlhG-GFP signal appears as discrete foci distributed along the cell body, consistent with flagellar insertion sites. Scale bar = 10 μm. (B) Demographic distribution of FlhG-GFP in *B. burgdorferi* cell poles (n=445 cells) and midcell (n= 269 cells).

### FlhG_Bb_ controls the polar localization of FlhF and FliF

FlhF is a key regulator of both the number and positioning of PFs in *B. burgdorferi* and is essential for maintaining the flat-ribbon flagellar architecture (13). Its function involves stabilizing critical flagellar components, including the MS-ring protein FliF. FliF was chosen because it forms the foundational MS-ring that nucleates assembly of the entire flagellar motor, making its proper localization critical for flagellar biogenesis (35, 46). To determine whether FlhG_Bb_ regulates flagellar patterning through FlhF and FliF, we expressed FlhF-GFP and FliF-GFP in their respective deletion mutants (*ΔflhF* and *ΔfliF*) and in the Δ*flhG* mutant.

Fluorescence microscopy revealed that FlhF-GFP localizes predominantly at the poles in the *ΔflhF* strain, with additional puncta at midcell during division (**Fig. 8A**). In contrast, deletion of *flhG_Bb_*completely abolished polar FlhF localization, resulting in dispersed puncta throughout the cell (**Fig. 8B**). Similarly, FliF-GFP formed distinct polar foci in the *ΔfliF* strain (**Fig. 9A**) but became diffused along the cell length in the Δ*flhG* mutant (**Fig. 9B**). Together, these results demonstrate that FlhG_Bb_ is essential for the proper subcellular localization of both FlhF and FliF, coordinating basal body assembly, flagellar positioning, and number.

**Figure 8.**
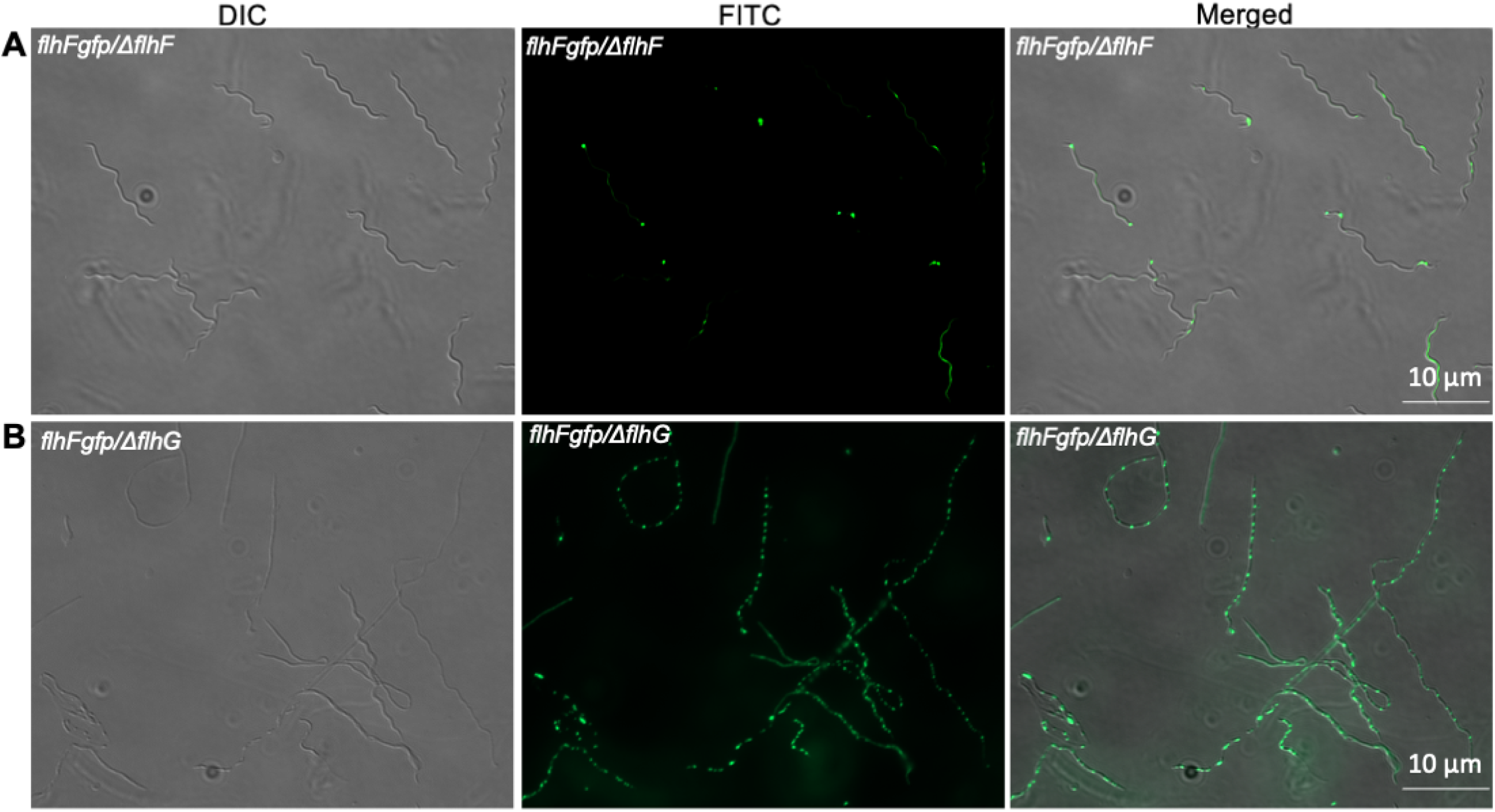
FlhGBb is required for the polar localization of FlhF. **(A)** Fluorescence microscopy of *flhFgfp/ΔflhF* cells expressing FlhF-GFP in a *ΔflhF* background. DIC (left), FITC fluorescence (middle), and merged (right) channels are shown. FlhF-GFP localizes as discrete polar or subpolar foci, consistent with flagellar tip/insertion localization. Scale bar = 10 μm. **(B)** Fluorescence microscopy of *flhFgfp/ΔflhG* cells expressing FlhF-GFP in a *ΔflhG* background. In the absence of FlhG, FlhF-GFP signal is redistributed along the cell body as multiple, evenly spaced foci rather than discrete polar clusters, consistent with derepressed flagellar assembly at ectopic sites. Scale bar = 10 μm.

**Figure 9.**
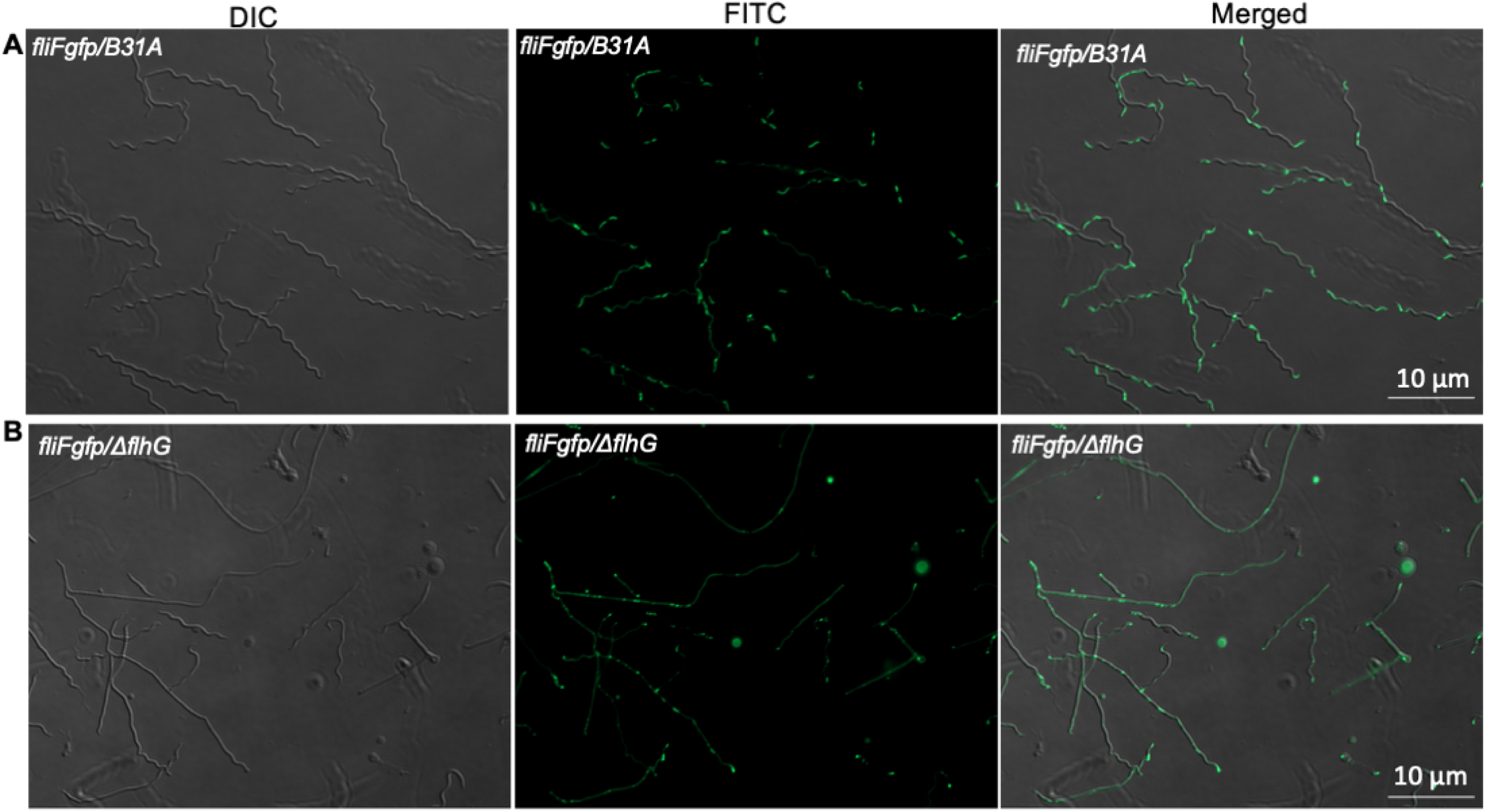
Loss of FlhG_Bb_ disrupts the polar localization of FliF. **(A)** Fluorescence microscopy of *fliFgfp*/B31A WT cells expressing FliF-GFP. DIC (left), FITC fluorescence (middle), and merged (right) channels are shown. FliF-GFP localizes as discrete foci at the cell poles, consistent with a limited number of polar flagellar motors per cell. Scale bar = 10 μm. **(B)** Fluorescence microscopy of *fliFgfp*/Δ*flhG* cells. In the absence of FlhG_Bb_, FliF-GFP foci are no longer restricted to the cell poles but are instead distributed along the entire cell length, indicating uncontrolled flagellar MS-ring assembly at ectopic sites. Scale bar = 10 μm.

## DISCUSSION

This study identifies FlhG_Bb_ (BB0269) as a spatial regulator that coordinates flagellar assembly, cell morphogenesis, and cytokinesis in *B. burgdorferi*. Although annotated as a FlhG-like ATPase, FlhG_Bb_ displays several structural features that distinguish it from canonical FlhG/MinD homologs, including a ∼30-residue N-terminal α-helical extension (**Fig. 1C**) and a repositioned interfacial helix (**Fig. 1E**). These structural differences likely reflect adaptations to the specialized cellular architecture of spirochetes. Consistent with its predicted classification within the MinD/ParA family of P-loop NTPases, FlhG_Bb_ is an active ATPase (**Fig. 1G**), and ATP hydrolysis is essential for its regulatory functions. Together, these findings establish FlhG_Bb_ as an energy-dependent spatial regulator that plays a pivotal role in organizing cellular architecture in *B. burgdorferi*.

A central insight from this work is that FlhG_Bb_ regulates flagellar biogenesis primarily through spatial organization rather than changes in flagellar protein abundance. A similar mode of regulation has been reported in *Campylobacter jejuni*, where FlhG does not control flagellar number by altering flagellar gene expression or protein production (33). In contrast, in many rod-shaped bacteria, FlhG homologs regulate flagellar number through transcriptional or post-translational pathways (16, 18, 24). For example, in *Vibrio* species, FlhG interacts with the transcriptional activator FlrA to repress flagellar gene expression and thereby limit polar flagellation (31). Similarly, the *Pseudomonas* FlhG homolog FleN controls flagellar number by modulating transcriptional regulators of the flagellar gene hierarchy. In spirochetes, deletion of *flhG* in *L. biflexa* has no impact on flagellar gene expression but instead alters the global transcriptome (14). Collectively, these observations suggest that although the role of FlhG homologs in regulating flagellation is broadly conserved, the underlying regulatory mechanisms vary across bacterial species. Consistent with this idea, our findings indicate that FlhG_Bb_ acts primarily at the level of subcellular organization, directing the localization of key flagellar assembly factors.

Specifically, FlhG_Bb_ controls the positioning of FlhF (**Fig. 8**), an SRP-type GTPase that specifies flagellar placement (13), and FliF, the MS-ring protein that nucleates basal body assembly (35, 47). Correct localization of these components ensures that basal bodies assemble at defined polar sites and generate the ribbon-like periplasmic flagellar bundles characteristic of spirochetes. When FlhG_Bb_ is absent, both FlhF and FliF lose polar confinement and appear as dispersed puncta along the cell body (**Figs. 8 & 9**), indicating that spatial control of early assembly events is critical for establishing the ordered architecture of the spirochete flagellar system. This regulatory mode resembles aspects of FlhG function described in *C. jejuni*, where FlhG influences polar flagellar placement (18, 33). However, the extensive spatial control observed here may be particularly important in spirochetes, whose PFs are mechanically integrated with the cell cylinder.

Our results further suggest that FlhG_Bb_ links flagellar assembly to the bacterial cell cycle. FlhG_Bb_ displays dynamic localization, concentrating at the poles in nascent cells and redistributing to midcell during division (**Fig. 7**). This pattern resembles the spatial behavior of MinD in rod-shaped bacteria (25, 45) and suggests that FlhG_Bb_ may provide positional cues that coordinate morphogenesis with cytokinesis. In this model, FlhG_Bb_ first promotes polar flagellar assembly by positioning FlhF and FliF, thereby defining sites for basal body formation. As the cell cycle progresses, redistribution of FlhG_Bb_ toward the division site may help coordinate septum formation with ongoing morphogenetic processes. Such coordination would be particularly important in spirochetes, where PF bundles span much of the cell length and could potentially interfere with septation if not properly positioned.

Beyond flagellar motor placement, FlhG_Bb_ also appears to influence higher-order flagellar motor architecture. Structural analyses revealed that the ring-like structure associated with the flagellar collar is absent in *ΔflhG* cells (**Fig. 5**). The collar is a distinctive feature that stabilizes the motor and anchors PFs within the cell envelope (40, 41). The absence of this structure in the mutant suggests that spatial cues provided by FlhG_Bb_ may influence later stages of motor assembly or stabilization. Thus, FlhG_Bb_ may coordinate multiple levels of flagellar morphogenesis, from the positioning of basal bodies to the assembly of specialized motor structures.

Together, these observations support a mechanistic model in which FlhG_Bb_ acts as a master spatial organizer in *B. burgdorferi* (**Fig. 10**). At the poles, FlhG_Bb_ directs the localization of FlhF and FliF to establish sites of basal body assembly and promote formation of organized PF ribbon bundles. As cells progress through the division cycle, redistribution of FlhG_Bb_ toward midcell likely contributes to proper septum placement, synchronizing cytokinesis with ongoing flagellar assembly. Through this ATPase-driven localization cycle, FlhG_Bb_ integrates morphogenesis, motility, and cell division to maintain the distinctive spiral morphology and highly ordered flagellar architecture of spirochetes.

**Fig. 10.**
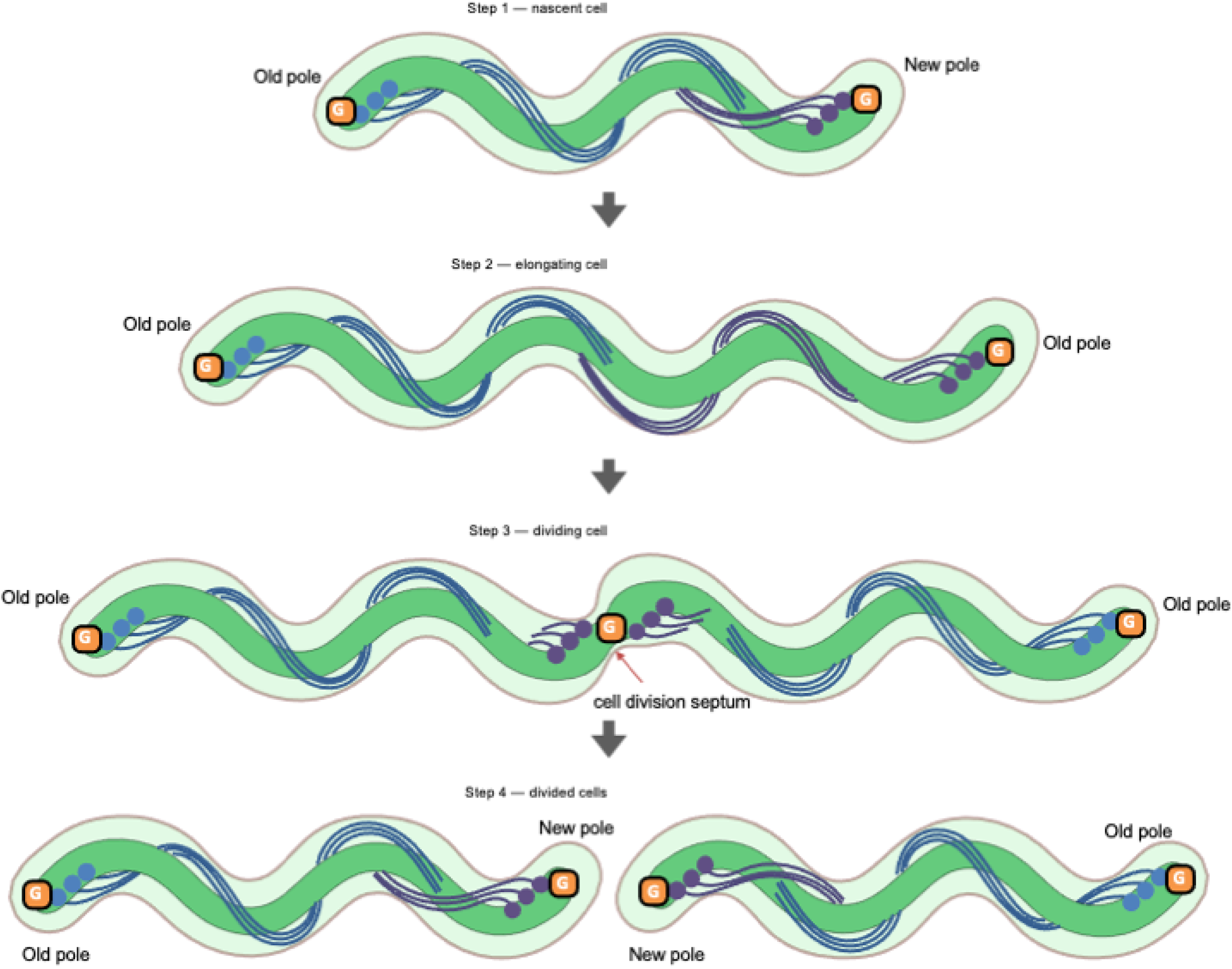
Model for FlhG_Bb_-mediated coordination of flagellation and cell division in *B. burgdorferi*. In nascent cells, the ATPase FlhG_Bb_ localizes predominantly at the cell poles, where it recruits and spatially organizes FlhF and the MS-ring protein FliF to establish sites of basal body assembly. These events initiate the formation of polar PFs that assemble into ribbon-like bundles along the cell cylinder. As the cell cycle progresses, FlhG_Bb_ redistributes toward the midcell division zone, where it likely contributes to proper septum placement and coordination of cytokinesis with ongoing flagellar morphogenesis. In the absence of FlhG_Bb_, FlhF and FliF become delocalized, basal bodies assemble at aberrant positions, and organized PF ribbon formation is disrupted, leading to defects in motility, cell morphology, and division. Through this ATPase-dependent spatial regulation, FlhG_Bb_ synchronizes flagellar assembly with cell division to maintain the distinctive architecture and motility of spirochetes.

More broadly, this work reveals how a conserved MinD-like ATPase has been functionally adapted to the specialized cell biology of spirochetes. Whereas canonical FlhG proteins primarily regulate flagellar number through transcriptional control pathways, FlhG_Bb_ appears to operate predominantly through ATPase-driven spatial organization of morphogenetic processes. This specialization likely reflects the unique structural constraints imposed by PFs, which must be precisely positioned and coordinated with cell growth to generate efficient motility. FlhG_Bb_ therefore represents a distinct paradigm for how conserved molecular scaffolds can evolve to integrate spatial organization, motility, and cell division in bacteria with specialized cellular architectures.

## Materials and Methods

### Bacterial strains and growth conditions

A high-passage *Borrelia burgdorferi* sensu stricto strain B31A (wild type) and its isogenic mutants were cultured in Barbour-Stoenner-Kelly II (BSK-II) liquid medium or on semi-solid agar plates at 34°C under 3.4% CO₂, as previously described(13, 48). When required, antibiotics were added for selective pressure at the following concentrations: kanamycin (300 μg/ml), gentamicin (40 μg/ml), or streptomycin (50 μg/ml). *Escherichia coli* NEB 5α (NEB, Ipswich, MA) was used for DNA cloning and plasmid propagation; M15 (Qiagen, Valencia, CA, USA) was used for recombinant protein expression. For *E. coli* selection, antibiotics were used at the following concentrations: kanamycin (50 μg/ml), spectinomycin (50 μg/ml), or ampicillin (100 μg/ml).

### Preparation of *B. burgdorferi* FlhG (FlhG_Bb_) recombinant protein and antibody

The gene (*BB0269*) encoding the full-length FlhG_Bb_ was PCR amplified using primers P_3_/P_4_ with engineered BamHI and PstI restriction sites at the 5′ and 3′ ends, respectively. The resulting amplicon was cloned into the pGEM-T Easy vector (Promega, Madison, WI) and then subcloned into the pQE80L expression vector (Qiagen, Valencia, CA), which encodes an N-terminal hexa-histidine (His₆) tag. The recombinant plasmid was transformed into M15 cells for protein expression. FlhG_Bb_ expression was induced with 1 mM isopropyl-β-D-thiogalactoside (IPTG). The His₆-tagged FlhG_Bb_ recombinant protein (His₆-FlhG_Bb_) was purified under denaturing conditions using a nickel-nitrilotriacetic acid (Ni-NTA) agarose column (Qiagen) and subsequently dialyzed overnight at 4°C against 10 mM Tris-HCl buffer. For antibody production, two rats were immunized with 1 mg of purified His₆-FlhG_Bb_ over a 1-month period, followed by booster injections (100 μg per rat) at weeks 6 and 7. Antiserum was generated by General Bioscience Corporation (Brisbane, CA).

### Site-directed mutagenesis

Site-directed mutagenesis was performed using the Q5 Site-Directed Mutagenesis Kit (NEB, Carlsbad, CA) according to the manufacturer’s instructions. The above constructed FlhG_Bb_ expression plasmid was used as the template. A single amino acid substitution (D69A) was introduced into FlhG_Bb_ using primer pair P_11_/P_12_. The mutation was confirmed by DNA sequencing. The mutated gene was subsequently PCR amplified and subcloned into the pQE80 expression vector (Qiagen) to generate an N-terminal His-tagged fusion protein.

### ATPase hydrolysis assays

Expression vectors encoding His₆-FlhG_Bb_ and its variants were transformed into *E. coli* strain Lemo21 (NEB, Ipswich, MA). Protein expression was induced with 1 mM IPTG for 18 h at 16 °C. Cells were harvested for protein purification using Ni-NTA agarose columns under native conditions. Protein concentrations were determined by Bradford assay. ATPase activity was measured by quantifying inorganic phosphate (Pi) release using a colorimetric ATPase/GTPase Activity Assay Kit (Sigma-Aldrich, St. Louis, MO), according to the manufacturer’s protocol. A standard curve generated with the supplied phosphate standard was used to convert absorbance at 620 nm to Pi concentration. For each reaction, 10 µL of purified protein (or assay buffer as a blank control) was mixed with 10 µL ATP and 20 µL assay buffer in a 96-well microplate. After incubation for 30 min at room temperature, 200 µL of colorimetric reagent was added, and the plate was incubated for an additional 30 min to allow color development. Absorbance at 620 nm was measured using a Variskan Lux microplate reader (Thermo Fisher Scientific, Waltham, MA).

For kinetic analysis, 80 nM purified His₆-FlhG_Bb_ was incubated with increasing ATP concentrations (ranging from 0 to 500 µM as indicated). Initial velocities were calculated from Pi production over time, and saturation curves were fit to the Michaelis-Menten equation using GraphPad Prism 10 (GraphPad Software, San Diego, CA). All assays were performed in triplicate in three independent experiments.

### FlhG model preparation and structural comparison

ATP-bound dimeric FlhG_Bb_ (Uniprot ID:Q44911) model was generated using AlphaFold3(49) (ipTM = 0.88, pTM = 0.90) and compared to AMPPNP bounds *Pseudomonas aeruginosa* FleN (PDB: 5J1J(26)) and *Escherichia coli* MinD (PDB: 3R9I(50)). Structures were visualized, analyzed and figures were prepared using PyMol.

### Construction of a *flhG* deletion mutant and its isogenic complemented strains

The vector for in-frame deletion of *flhG* (*BB0269*) was generated using a PCR-based fusion strategy as previously described(13). Briefly, primer pairs P_5_/P_6_ and P_7_/P_8_ (the latter containing sequences complementary to *aadA1*, a streptomycin resistance marker) were designed to amplify approximately 1-kb regions upstream and downstream of the *flhG* sequence. Three separate PCR reactions were performed to amplify the 5′- and 3′-flanking regions of *flhG* and the *aadA1* cassette(51). The resulting PCR fragments were fused by overlap-extension PCR using primers P_5_/P_8_, generating the deletion construct (*flhG*::aadA1). The resulting construct was transformed into wild type competent cells by electroporation. Mutant clones were selected on semi-solid agar plates supplemented with streptomycin (50 μg/ml). The resulting mutant clones were confirmed by PCR and DNA sequencing. One mutant clone (*ΔflhG*) was chosen for cis-complementation through inserting the full-length *flhG_Bb_* gene into an intergenic region between *BB0445* and *BB0446* on the chromosome, as previously described (30, 52). Primer sequences used here are listed in **Table S1**.

### Measurement of *B. burgdorferi* growth rates

To determine growth kinetics, early stationary-phase cultures of wild-type (WT) B31A, the *flhG* deletion mutant (Δ*flhG*), and the cis-complemented strain (Δ*flhG*^com^) were diluted into fresh BSK-II medium to a final density of 1 × 10⁵ cells/ml and incubated at 34°C. Cell densities were measured daily for up to 8 days using a Petroff–Hausser counting chamber(30). Counts were performed in triplicate for three independent cultures. Data are presented as means ± standard errors of the mean (SEM).

### Bacterial motion tracking analysis and swimming plate assays

The swimming velocity of *B. burgdorferi* cells was quantified using a computer-based motion-tracking system as previously described(8, 12, 53). Briefly, log-phase bacterial cultures were diluted 1:1 in BSK-II medium. A 20-μl aliquot of the diluted culture was mixed with an equal volume of 2% methylcellulose (MC4000; viscosity, 4,000 cP). Cells were recorded using iMovie software on an Apple Mac computer. Videos were exported as QuickTime files and imported into OpenLab (Improvision Inc., Coventry, UK), where frames were cropped, calibrated, and saved as LIFF files. Individual cells were tracked and velocities calculated using Volocity software (Improvision Inc., Coventry, UK). For each strain, at least 20 cells were analyzed for up to 30 s. Swimming plate assays were performed using 0.35% agarose prepared with BSK-II medium diluted 1:10 in Dulbecco’s phosphate-buffered saline (DPBS; pH 7.5), as previously described(12, 48). Swimming ring diameters were measured in millimeters (mm), and mean values were calculated from four independent plates for each strain.

### Gel electrophoresis and Western blot analysis

Sodium dodecyl sulfate-polyacrylamide gel electrophoresis (SDS-PAGE) and immunoblotting using enhanced chemiluminescence (ECL) detection were performed as previously described(13, 37). Briefly, 10-20 μg of total cell lysate was resolved on 12% SDS-PAGE gels and transferred to polyvinylidene difluoride (PVDF) membranes (Bio-Rad Laboratories). Membranes were probed with specific primary antibodies against *B. burgdorferi* proteins, followed by horseradish peroxidase (HRP)-conjugated secondary antibodies and detection using an ECL luminol-based assay. Monoclonal anti-FlaB (H9724) was kindly provided by A. G. Barbour (University of California, Irvine, CA), and monoclonal anti-DnaK by J. Benach (SUNY at Stony Brook, NY). Polyclonal antibodies against FlgG, FliF, FlgE, FlhF, and FliD were described previously(13, 37, 54). Chemiluminescent signals were captured using a ChemiDoc MP imaging system and quantified with Image Lab software (Bio-Rad Laboratories, Hercules, CA). Three independent biological replicates were analyzed for each experiment.

### Construction of plasmids that express FlhG_Bb_-GFP and FliF_Bb_-GFP fusion proteins

These vectors were constructed as previously documented(13). To make the constructs, the full-length *flhG_Bb_* and *fliF_Bb_*genes were PCR amplified with primer sets P_9_/P_10_ and P_13_/P_14_, respectively, in which XhoI and SacII cut sites were engineered. The resultant PCR fragment was first cloned into pGEM-T vector and then subcloned into precut *P_flgB_*-*flhF_Bb_*-*gfp*/V2G at the site of XhoI and SacII, yielding the vector of *P_flgB_*-*flhG_Bb_*-*gfp*/V2G or *P_flgB_*-*fliF_Bb_*-*gfp*/V2G. The final construct was confirmed by DNA sequencing.

### Light and fluorescence microscopy

Cell morphology and motility of *B. burgdorferi* strains were characterized using light microscopy as previously described(13, 55, 56). In brief, *B. burgdorferi* cells in BSK-II medium were videotaped with dark-field illumination at 100× for at least 30 seconds. Videos were digitized to allow for frame-by-frame observation of cell motion. Fluorescence images were captured using a Zeiss Observer D1 Inverted LED fluorescence microscope at a wavelength of 480 nm using a Zeiss Observer D1 inverted LED fluorescence microscope equipped with an excitation filter (541-569 nm) and an emission filter (581-654 nm). The images were processed using the ZEN Blue software (Zeiss, Germany).

### Fluorescent D-amino acid labeling

HADA (3-[7-hydroxycoumarin]-carboxamide-D-Alanine) were synthesized by Tocris following the published protocol(43). Mid-log phase *B. burgdorferi* cultures (2∼3×10⁷ cells/mL) grown in BSK-II medium were pelleted (9,000 × g, 10 min, RT), resuspended in fresh medium, and incubated with 100µM HADA for 60-120 min at 34°C. Cells were briefly pelleted (9,000 × g, 5 min), washed three times with PBS (pH 7.4), and resuspended in PBS for live imaging. Fluorescence images were taken using a Zeiss Observer microscope with a DAPI/FITC filter set. The images were captured and processed using the program ZEN Blue (Zeiss, Germany). For demography analysis, Fiji ImageJ software was used to generate data frame for each cell. At least 400 cells were analyzed per condition. The resulting data were organized into demographs using NumPy(57) and plotted with Matplotlib(58).

### Sample preparation for cryo-electron tomography (cryo-ET), imaging, and data processing

*B. burgdorferi* cells in liquid culture were resuspended in PBS to a final OD₆₀₀ of 1.0. A 10-nm gold tracer solution (Aurion) was added, and 5 μL of the mixture was applied to glow-discharged cryo-EM grids (Quantifoil). The grids were blotted from the front with Whatman™ filter paper and then plunged into a liquid ethane/propane mixture using a Leica EM GP2 plunger. The GP2 environmental chamber was maintained at 25°C and 95% humidity, and grids were blotted from the front for 6 s.

Frozen-hydrated specimens were imaged at below -180°C using a Titan Krios G2 300 kV transmission electron microscope (ThermoFisher) equipped with a field emission gun, K3 direct detection camera (Gatan), and GIF BioQuantum K3 Imaging Filter (Gatan). Tilt series were recorded with SerialEM software (59) at 42,000× magnification, with a physical pixel size of 2.148 Å and a defocus of 4.8 μm. The stage was tilted from -48° to +48° in 3° increments using a dose-symmetric scheme in the FastTomo script (DOI: 10.1101/2021.03.16.435675). The total electron dose was ∼70 e⁻/Å² distributed over 33 images per tilt series, recorded in dose-fractionation mode.

MotionCor2 (60) was used to correct image drifting induced by the electron beam during image recording. IMOD software was used to create image stacks and then track with fiducial beads to align all images in each tilt series (61, 62). Gctf (63) was used to estimate defocus for all aligned images, and the contrast transfer function (CTF) was corrected using the ctfphaseflip function in IMOD(64). To reconstruct 8x binned tomograms, 8x binning of images in the tilt series was created using the binvol function in IMOD. Tomo3D (65) was then used to reconstruct tomograms. Collectively, 148 tomograms of Δ*flhG* mutant cells and 129 tomograms of Δ*flhG* complementary cells were collected (see details in **Table S2**). The software package Dragonfly (Version 2022.2, Comet Technologies Canada Inc.) was utilized to segment the bacterial features, including the outer membrane, inner membrane, and flagellar filaments of varying lengths.

### Statistical analysis

For growth curves, swimming plate assays and bacterial cell motion tracking analysis, the results are expressed as means ± standard errors of the means (SEM). The significance of the difference between different experimental groups was evaluated using ANOVA (*P* value < 0.01).

## Acknowledgement

This work was supported by funding from the National Institutes of Allergy and Infectious Diseases (AI078958 to C. Li; AI087946 to J. Liu; GM122535 to B. Crane; and AI148844 to B. Crane and C. Li), National Institutes of Health (NIH). Cryo-ET data were collected at Yale Cryo-EM Resource, which was funded in part by the NIH grant 1S10OD023603-01A1. We thank the Yale Center for Research Computing (YCRC) for guidance and use of the research computing infrastructure and Dr. Joshua W McCausland (Department of Biology, Stanford University) for providing us codes for demographic analysis.

## Author contributions

K.Z., W.G. and M.J.L performed the experiments, analyzed the data, and wrote the manuscript. B.R.C, J.L. and C.H.L. secured the funding and contributed to the design and supervision of the study and the revision of the manuscript. All authors read and approved the final version of the manuscript.

## Movie legends

Movie 1: Video tracking analysis of the wild type strain B31A.

Movie 2: Video tracking analysis of the *flhG* deletion mutant (*ΔflhG*).

Movie 3: Video tracking analysis of the complemented strain (*ΔflhG^com^*).

Movie 4: Video tracking analysis of FlhG D69A point mutant (*ΔflhG^D69A^*).

